# MUC5B mobilizes and MUC5AC spatially aligns mucociliary transport on human airway epithelium

**DOI:** 10.1101/2022.03.11.484020

**Authors:** Daniel Song, Ethan Iverson, Logan Kaler, Margaret A. Scull, Gregg A. Duncan

## Abstract

Airway mucus acts as a protective barrier and vehicle for clearance of pathogens, providing the lungs with a defense mechanism called mucociliary clearance (MCC). Airway mucus is composed of two mucins, mucin 5B (MUC5B) and 5AC (MUC5AC) that form a hydrogel that enables functional clearance in health. However, abnormalities in mucin expression, specifically increases in MUC5AC is observed in chronic respiratory diseases and leading to defective MCC. Our current understanding of MCC impairment focuses on mucin concentration, while the impact of mucin composition remains unclear. Here, we use MUC5AC/B-knock out (KO) human airway epithelial (HAE) tissue cultures to understand the role and contribution of individual secreted mucins on MCC mechanisms. We find that KO cultures result in impaired or discoordinated mucociliary transport demonstrating the importance of each of these mucins to effective MCC and shedding light on a new mechanism of mucin composition-dependent airway clearance.

## Introduction

Airway mucus is a complex biological fluid that protects the lung from environmental insults through a process known as mucociliary clearance (MCC). The mucus layer, which coats the airways, protects the underlying epithelium by acting as a physicochemical barrier immobilizing inhaled pathogens and particulates (*1*, *2*). Unlike other mucosal tissues, the airways can dynamically clear mucus through shear forces exerted by the rapid beating of cilia on the epithelium allowing removal of trapped particulates (*3*, *4*). Normal MCC is essential for innate defense and is facilitated by a transportable mucus layer and coordinated ciliary activity (*5*).

The primary secreted mucins expressed in the lung are mucin 5B (MUC5B) and 5AC (MUC5AC) (*6*, *7*). These mucins are large (~MDa) glycoproteins that have cysteine-rich domains at their N- and C-termini that cross-link and polymerize via disulfide bonds (*4*, *8*). The resulting network acts as a barrier by controlling the transport of nano- and microscale entities through physical and steric interactions while also giving rise to the viscoelastic nature of mucus (*9*–*13*). While MUC5B and MUC5AC have a high degree of similarity in amino acid sequence and domain organization, recent evidence shows that MUC5B and MUC5AC differ in assembly and N-terminal organization, with MUC5B mainly forming dimers, and MUC5AC assembling into dimers as well as higher-order oligomers (*14*, *15*). These subtle differences in macromolecular organization impact the functional properties of secreted mucins, such as effectiveness as a barrier and transportability, which have been studied in *in vivo* animal models (*16*). In Muc5ac/b knockout (KO) mice, Muc5b deficiency leads to impaired mucus transport and pathogen accumulation, while Muc5ac is shown to be dispensable for transport (*17*, *18*). Nonetheless, in health, the two gel-forming mucins form a mucus gel that is optimized for providing proper protection to the airways.

In health, MUC5B is the predominant mucin, while MUC5AC is expressed at lower levels (*6*, *7*). While the expression of MUC5B and MUC5AC is tightly regulated to ensure efficient MCC, production of MUC5AC is increased in chronic respiratory diseases due to type I and II immune responses, resulting in a mucus gel with abnormal mucin composition (*7*, *19*–*21*). Elevated MUC5AC in mucus has been identified as the major driver in impairment of MCC and is correlated with enhanced viscoelastic properties, decreased lung function in chronic bronchitis, and significant mucous plugging in fatal asthma (*7*, *19*, *22*, *23*). However, how MUC5AC, in health, contributes to effective mucus clearance, while driving the formation of stagnant mucus in disease is unknown. As a result, there is growing interest in understanding how these two secreted mucins with similar domain structures but varying functional properties work in concert to enable efficient airway clearance. Moreover, the spatial coordination of transport on airway surfaces has been largely attributed to cilia organization and activity (*24*). However, the contribution of secreted mucins to tissue-scale alignment in MCC has not been widely considered.

In our studies, we developed *in vitro* airway epithelial systems deficient for one of the two gel-forming mucins to investigate the role and contribution of MUC5B and MUC5AC on MCC mechanisms. We used an *in vitro* human airway epithelium model to harvest mucus gels composed of single mucins for rheological assessment. Using our cultures, we measured mucociliary transport rates, while also assessing flow orientation in real-time, demonstrating that MUC5B deficiency leads to impaired mucus clearance, whereas MUC5AC deficiency leads to transport that lacks spatial coordination. Accordingly, we observed that that supplementation of KO cultures with the absent mucin could improve these irregular mucociliary transport phenotypes. Taken together, our data demonstrates the functional significance of both mucins for promoting effective MCC in health, while providing insight into the mechanism of composition-dependent MCC impairment.

## Results

### Generation of MUC5B/AC-KO BCi-NS1.1 cultures

We used CRISPR/Cas9-mediated genome editing to generate *in vitro* HAE models genetically deleted for one of the two gel-forming mucins. We designed two single guide (sg)RNAs that target each mucin gene, MUC5B and MUC5AC, at different exons (**Table S1**). Earlier exons were targeted to maximize knock-out efficiency, while two sgRNAs were designed to account for potential off-target effects. As our control, we used a sgRNA with no predicted target. The sgRNAs were cloned into a GFP-expressing lentiviral vector that also encodes the Cas9 nuclease. After lentivirus production, we transduced BCi-NS1.1 cells, an immortalized HAE cell line, which has previously been used for manipulating mucin expression (*25*–*27*). Subsequently, transduced cells were expanded, and sorted for GFP expression prior to differentiation (**Fig 1A**). Using a T7 endonuclease cleavage assay, we confirmed editing at the correct genomic loci in undifferentiated cells, evidenced by heteroduplex fragments in all KO conditions (**Fig 1B, C**). To verify that lentiviral transduction did not impact normal differentiation of BCi-NS1.1 cultures, we measured the transepithelial electrical resistance (TEER) 28 days into air-liquid interface (ALI) culture and confirmed TEER values that were within reported ranges in all conditions, indicative of cell polarization and differentiation into a pseudostratified epithelium (**Fig S1)** (*28*).

**Figure 1.**
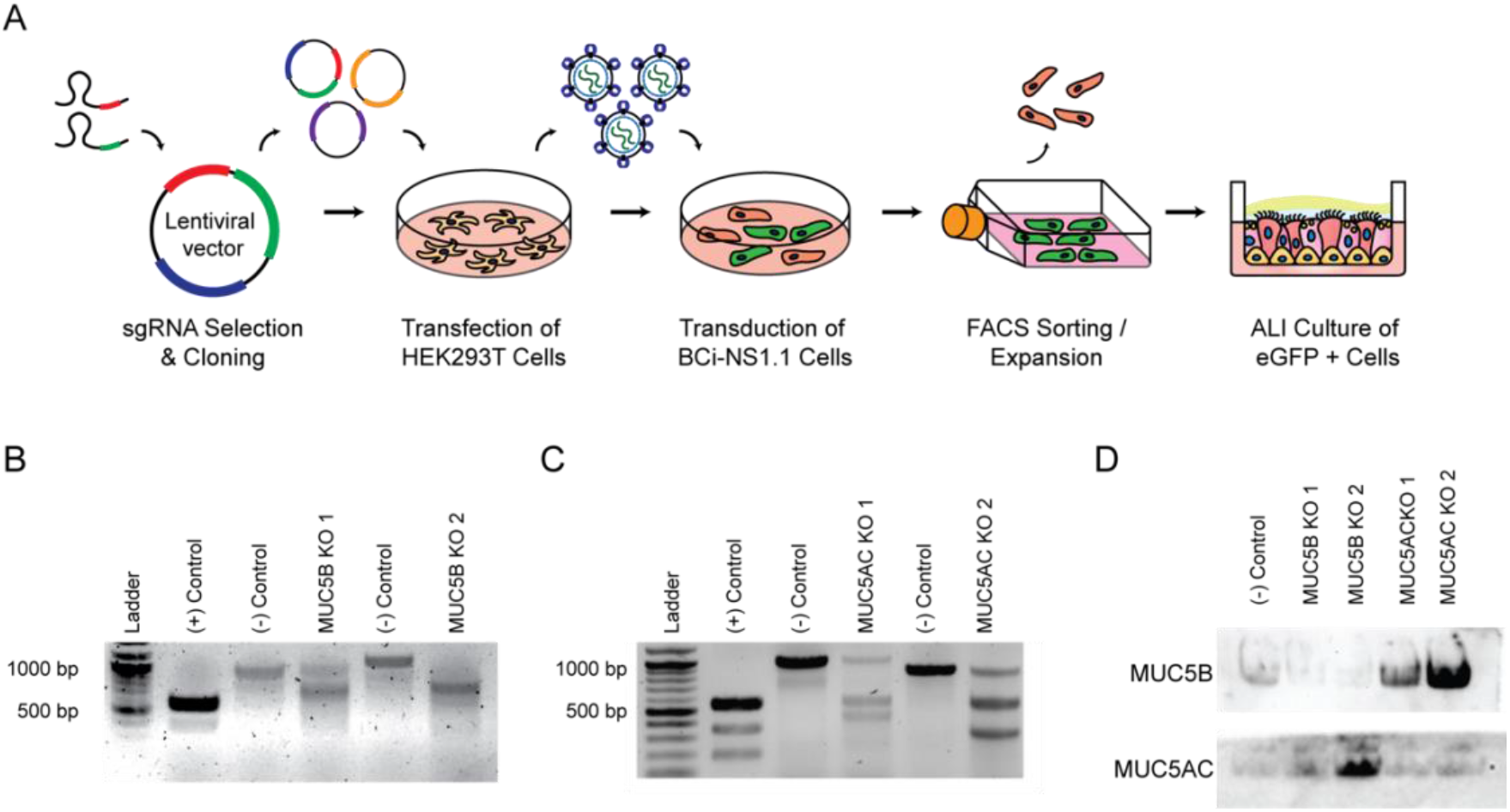
Generation and validation of MUC5B/AC-KO cultures. **(A)** Overview of MUC5B/MUC5AC gene targeting approach via lentivirus-mediated delivery of sgRNA and CRISPR/Cas9 machinery. (**B-C**) Validation of on-target editing using T7 endonuclease cleavage assay. Gel images show PCR products amplified from target sites where digested fragments indicate on-site editing. KO1 corresponds to BCi-NS1.1 cells transduced with sgRNA1 and KO2 corresponds to cells transduced with sgRNA2. (+) control corresponds to a control template and primer mix provided with the assay kit. (-) control corresponds to BCi-NS1.1 cells treated with a non-targeting sgRNA. (**B**) T7 endonuclease cleavage assay using genomic DNA from MUC5B-KO cultures. (**C**) T7 endonuclease cleavage assay using genomic DNA from MUC5AC-KO cultures. (**D**) Western blot analysis of mucus samples collected from MUC5B and MUC5AC-KO cultures.

Using mucus samples collected from apical washes, we confirmed KO expression on western blot analysis demonstrating loss of MUC5B and MUC5AC expression in KO cultures using respective sgRNA (**Fig 1D**). These data also confirmed that mucus secretions from MUC5B-KO cultures are primarily composed of MUC5AC, while secretions from MUC5AC-KO cultures are composed of MUC5B, allowing us to study the native properties of mucus composed of single mucins.

### Viscoelasticity of MUC5B and MUC5AC gels

To evaluate the biophysical properties of gels composed of either MUC5B and MUC5AC, we performed macro- and microrheology on pooled mucus samples collected from control and KO cultures. Using macrorheology, we observed elastic dominant properties for all mucus samples (G’>G’’), confirming a cross-linked polymer network. We also demonstrated that samples exhibited rheological properties in range of previously reported values for mucus collected *in vitro* and *ex vivo* (**Fig 2A**; **Fig S2**) (*11*–*13*). While MUC5AC gels resulted in slightly higher mean complex viscosities than other gel types, differences were not statistically significant (**Fig 2B**). We next measured percent solids concentration of each sample as the biophysical properties of a polymer gels are highly sensitive to changes in solids concentration. We found that the solids concentration of pooled mucus samples were within a physiological range (2-4%) (**Fig 2C**) (*4*, *11*). While macrorheological properties have major implications for flow and transport, mucus microstructural network allows mucus to act as a barrier to inhaled material by controlling the diffusion of nano- and microscale entities through physical and adhesive interactions (**Fig 2D**) (*4*). Therefore, we used particle tracking microhreology (PTM) to assess diffusion of muco-inert nanoparticles (MIP) through mucus with varying mucin composition to probe mucus microstructure. Results showed that MUC5AC gels had significantly reduced MIP diffusion, as measured by mean square displacement (MSD), and reduced pore size compared to MUC5B gels and mucus collected from control cultures (**Fig 2E, F**). Comparing the frequency distributions of MIP diffusion, we found that MUC5B gels resulted in MIP diffusion that was similar to mucus obtained from control cultures, whereas MUC5AC gels resulted in a significant reduction in MIP diffusion with multiple peaks indicative of mucus gels with a tighter mesh network and heterogenous structure (**Fig 2E, F, G, I**). Importantly, we also found that there were no significant differences in macro– and microrheology measurements of MUC5B and MUC5AC gels harvested from cultures with equivalent KO condition (KO1 and KO2) targeted using different sgRNAs.

**Fig. 2.**
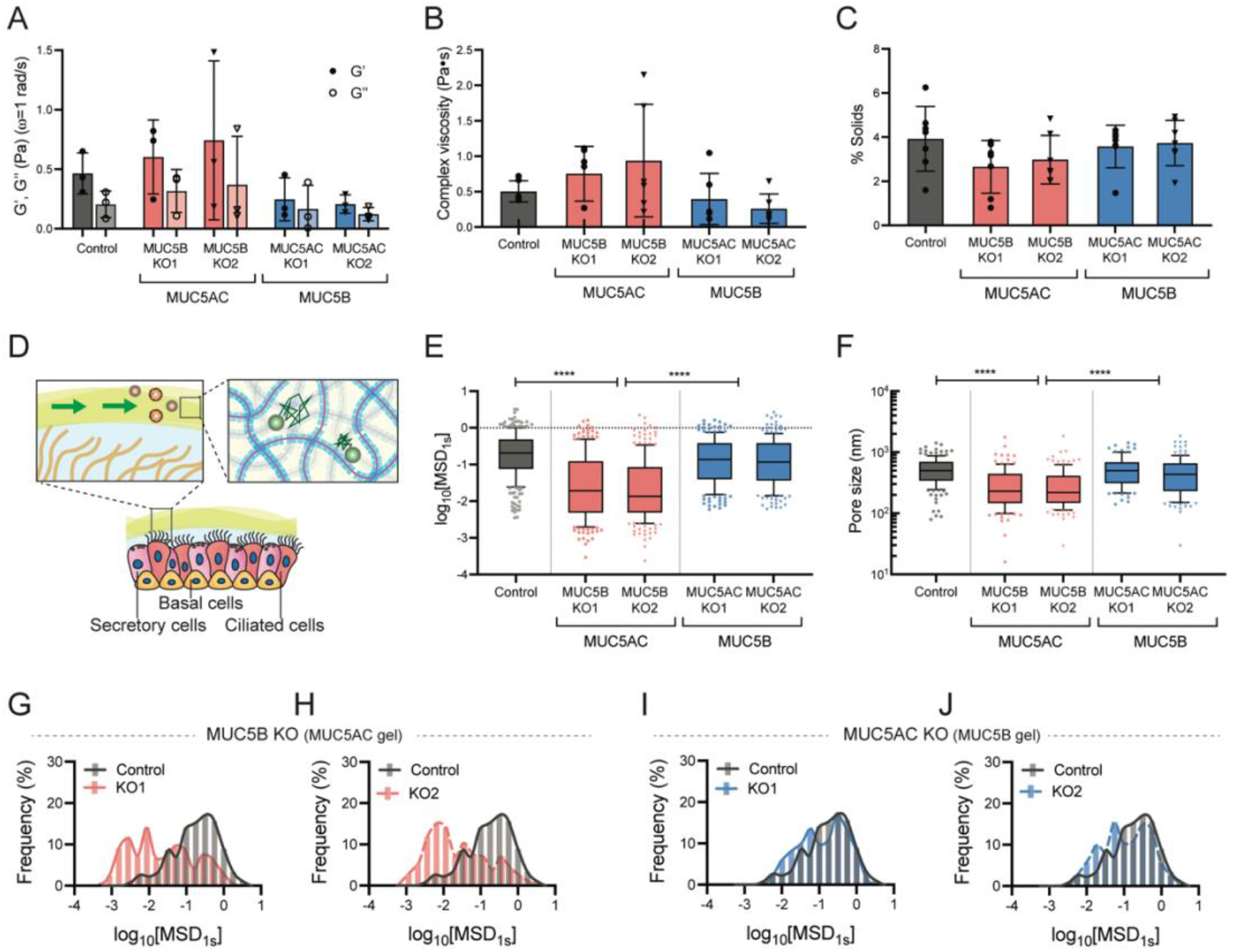
Viscoelastic properties of mucus produced from MUC5B/AC-KO cultures. (**A**) Elastic and viscous moduli (G’, G”) at ω = 1 rad/s measured using macrorheology. Moduli show G’-dominant mucus gels. (**B**) Complex viscosity measured using macrorheology. (**C**) Concentration of % solids in pooled mucus samples from MUC5B/AC-KO and control cultures. (**D**) Airway mucus acts as a vehicle of clearance of pathogens, providing the lungs with a robust defense mechanism. Mucus also has a microstructural network that can immobilize diffusing particles. (**E**) Diffusion of 100 nm PEG-NP in mucus samples from KO cultures as measured by logarithm mean squared displacement (MSD). (**D**) Estimated pore sizes of mucus samples. (**G-J**) Frequency distribution of log_10_[MSD_1s_]. Particle distribution in control and **(E**) MUC5B-KO1, **(F**) MUC5B-KO2, (**G**) MUC5AC-KO1, and (**H**) MUC5AC-KO2 samples.

### MUC5B-KO cultures exhibit reduced mucociliary transport and MUC5AC-KO cultures exhibit discoordinated transport

We next examined the mucociliary transport rates of mucus produced from MUC5B or MUC5AC KO cultures to investigate the contribution of each gel-forming mucin on mucociliary clearance. To do so, we deposited 2-μm microspheres onto KO cultures and tracked the movement of particles in real-time (**Fig 3A**). We observed that movement of microspheres on MUC5B-KO cultures were significantly reduced compared to the non-targeted control cultures. While functional transport was maintained in MUC5AC-KO cultures, we observed loss of unidirectional flow (**Fig 3B**). Quantifications of microsphere speed confirmed that MUC5B-KO cultures had significantly reduced transport rates compared to control and MUC5AC-KO cultures (**Fig 3C**). We next quantified the directionality of mucus flow on control and MUC5AC-KO cultures by measuring the angle, Δθ, which was calculated by measuring the angle between individual microsphere trajectory and ensemble average flow direction (θ_Average_) (**Fig 3D**). We found that control cultures resulted in an overall alignment of flow direction, where 95% of the microspheres moved within 10° to −10° of overall flow (**Fig. 3E**). However, MUC5AC-KO cultures resulted an irregular flow pattern that resembled circular swirls, where only 35-50% of microspheres aligned with the ensemble average flow direction (**Fig 3F,G**). We confirmed that these differences in transport rate and directionality of mucus flow were not likely the result of changes in the periciliary layer properties by confirming normal cilia beat frequency and normal cilia expression as demonstrated in +tubulin % area (**Fig 3H-J, Fig S3**).

**Figure 3.**
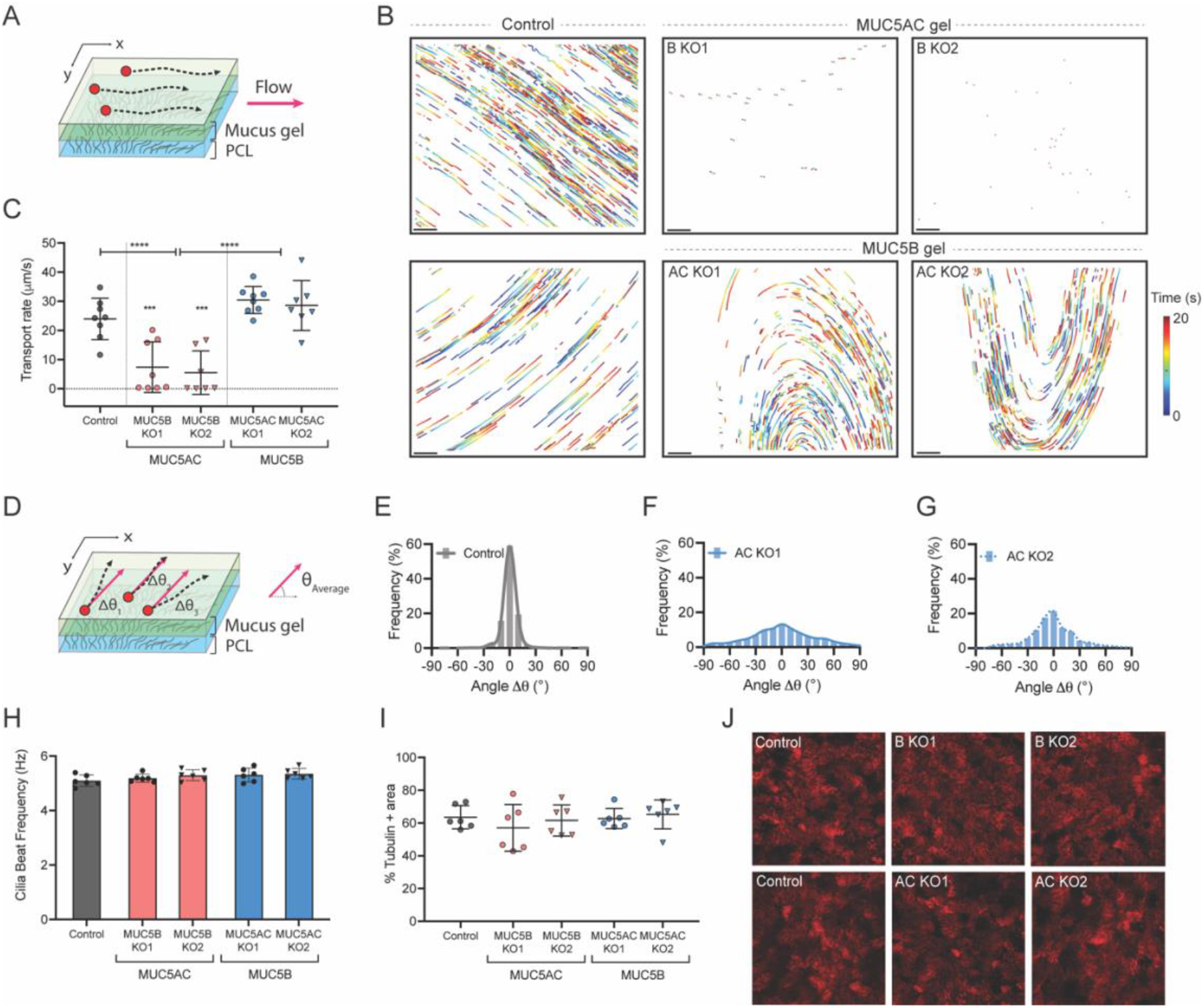
Mucociliary clearance dynamics of MUC5B/AC-KO cultures. **(A)** Mucociliary transport rate is measured by apically applying 2-μm microspheres to cultures and measuring particle displacement over time. **(B)** Representative trajectory images of microspheres over time. Trajectories show 20 s of motion with a color scale that indicates elapsed. Scale bar = 100 μm. (**C**) Mucociliary transport rates of MUC5B/AC KO cultures. (**D**) Trajectory angle, Δθ, was calculated by measuring the angle between and the overall flow direction (θ_Average_) and instantaneous individual microsphere trajectory. θ_Average_ was set to 0°. (**E-G**) Frequency distribution of Δθ. (**E**) Control (**F**) MUC5AC-KO1 (**G**) MUC5AC-KO2. (**H)** Cilia beat frequencies of MUC5B/AC KO cultures. (**I**) Quantification of +tubulin % area. (**J**) Representative image of cilia expression of MUC5B/AC-KO cultures by immunostaining of acetylated alpha tubulin.

### Transplanting exogenous mucus rescues impaired clearance

Our previous data showing abnormal transport resulting from loss of one of the two mucins led us to hypothesize that supplementing KO cultures with the absent mucin could restore stalled and/or irregular mucociliary transport phenotypes. To test this, we transplanted exogenous MUC5B and MUC5AC gels onto KO cultures and tracked the movement of microspheres on the cultures after equilibration (**Fig 4A**). We chose MUC5B-KO1 and MUC5AC-KO2 as representative cultures for transplantation experiments. We found that transplantation of MUC5B onto MUC5B-KO cultures resulted in a significant increase in transport rates, while also resulting in unidirectional flow (**Fig 4B-D**). However, transplantation of MUC5AC gels onto their native MUC5B-KO cultures did not result in a significant change in transport rate. Interestingly, the addition of either MUC5B or MUC5AC gels onto MUC5AC-KO cultures did not result in any significant change to transport rates (**Fig. 4E**) but resulted in improved directionality as shown in the trajectory images (**Fig 4F,G**).

**Figure 4.**
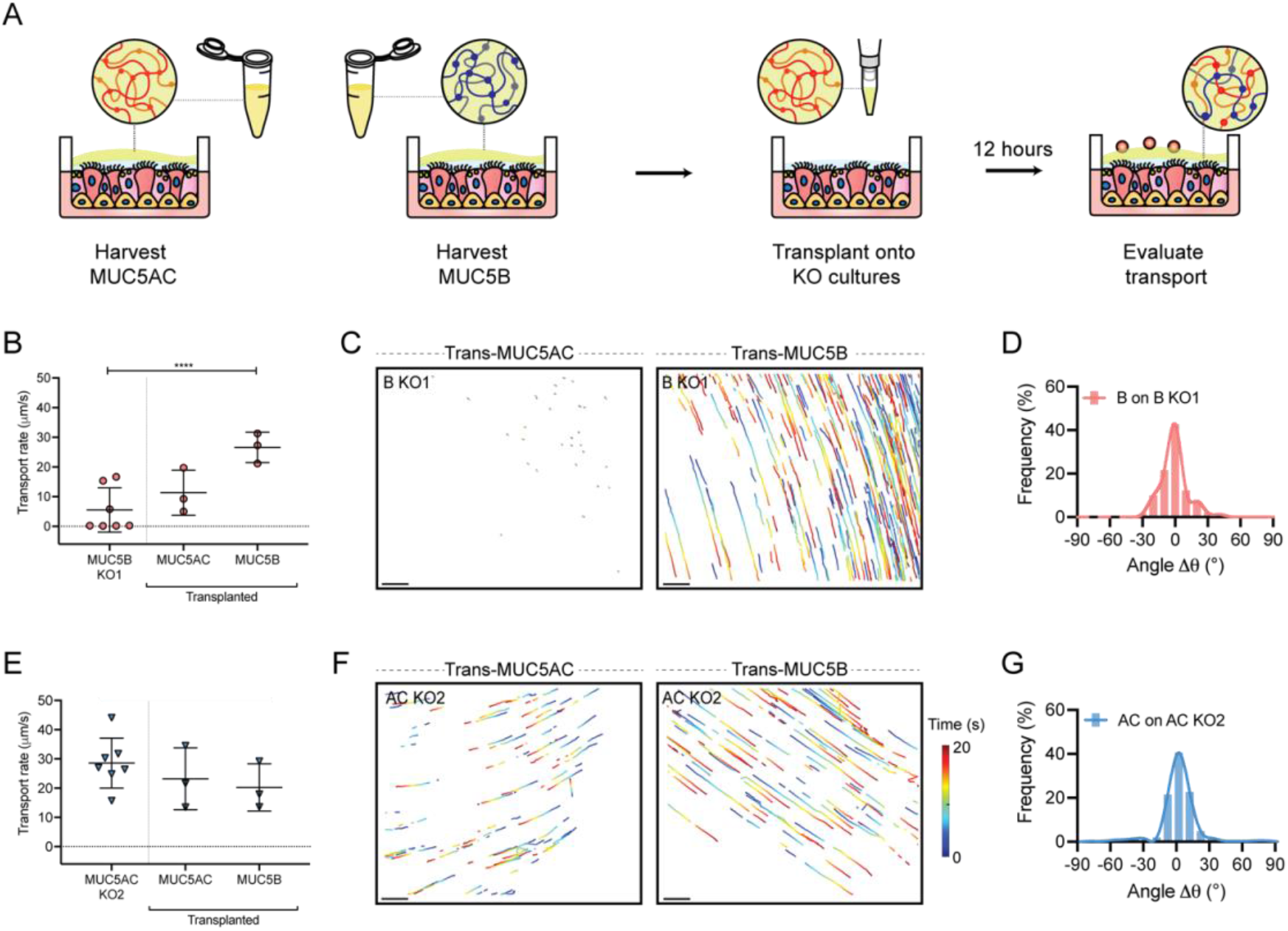
Mucociliary clearance dynamics of transplanted mucus on MUC5B/AC-KO cultures. **(A)** Exogenous mucus collected from KO cultures was transplanted onto KO cultures to supplement KO cultures deficient mucin. **(B)** Mucociliary transport rates of transplanted gels on B-KO culture (MUC5B-KO1 was used). (**C**) Representative trajectory images of microspheres trajectories on transplanted mucus on B-KO cultures. Scale bar = 100 μm. (**D**) Frequency distribution of Δθ, demonstrating directional flow. (**E)** Mucociliary transport rates of transplanted gels on AC-KO culture (MUC5AC-KO2 was used). (**C**) Representative trajectory images of microspheres trajectories on transplanted mucus on AC-KO cultures. Scale bar = 100 μm. (**D**) Frequency distribution of Δθ, demonstrating directional flow.

## Discussion

In these studies, we report *in vitro* HAE models that are genetically depleted for one of the two gel-forming mucins expressed in airway mucus to understand their role and overall contribution in mucociliary clearance mechanisms. We targeted MUC5B and MUC5AC genes in the BCi-NS1.1 human airway basal cell line and demonstrated successful editing at targeted genomic loci in undifferentiated cultures, while confirming KO expression in pooled mucus washings post-differentiation. MUC5B-KO cultures produced gels composed primarily of MUC5AC, while MUC5AC-KO cultures produced MUC5B gels, which allowed us to study each mucin independently. We note that our western blot analysis indicates that KO cultures produce slightly elevated levels of respective mucins compared to control cultures, while overall percent solids and protein concentration are conserved. These results may indicate that MUC5B and MUC5AC secretion is hydrodynamically controlled, regardless of mucin type. However, the pathways that regulate secretion of each mucin after KO will need to be explored in future studies.

Using our KO cultures, we assessed the biophysical properties of mucus samples with varying mucin composition. Our results indicate that gels composed of MUC5AC have altered microstructural properties compared to MUC5B gels, where pore size is significantly reduced and heterogenous. These results are consistent with recent findings that show that MUC5AC secreted from CALU3 cells form tight networks with high degrees of branching, while MUC5B organizes in linear strands (*15*). The smaller pore size in MUC5AC gels have implications for their functional role as a protective gel against pathological and chemical insults to the airway. In past work, MUC5AC has been shown to protect against viral respiratory pathogens such as influenza and prevent parasite infiltration (*21*, *29*, *30*). We also observed major differences in transportability of gels composed of different mucin types. MUC5AC gels produced from MUC5B-KO cultures were poorly cleared, while MUC5B gels from MUC5AC-KO cultures were more easily transported. These data are consistent with previous mouse models that showed that Muc5b-KO in mice, but not Muc5ac-KO, resulted in impaired mucociliary clearance and mucus accumulation in the airways (*17*). Furthermore, in humans, new evidence has shown that congenital absence of MUC5B results in impaired mucus clearance (*31*). This may be due in part to macromolecular organization of mucins, where MUC5B forms linear gels that can be easily transported via ciliary activity, while MUC5AC forms a tighter gel, which increases friction between gel surfaces, resulting in poor clearance (*3*, *15*). While bulk rheological properties are known to impact transport properties of the mucus gels, our data showed no resolvable differences in viscosities between gels of varying mucin types (*4*). Previous studies have predicted MUC5AC to form more elastic gels than MUC5B and we note that MUC5AC gels had higher complex viscosities compared to MUC5B gels, despite having lower percent solids concentration (*15*). We expect solids concentration-matched MUC5B and MUC5AC gels would possess even greater differences in their viscoelastic properties and this will be explored in subsequent work.

Despite functional transport of MUC5B gels on MUC5AC-KO cultures, we observed that MUC5AC deficiency led to poor directionality with local swirls, while control cultures that expressed both mucins showed global order. These transport patterns have been previously observed in HAE *in vitro* models and attributed to hydrodynamic active coupling between mucus and cilia, where changes in mucus properties and cilia density can alter the overall orientation of mucus flow (*32*, *33*). While the role of MUC5AC in mucociliary transport has been largely overlooked, our data suggests that MUC5AC–enhanced mucus viscoelasticity plays a significant role in hydrodynamic coupling between the mucus and periciliary layers, driving long-range transport and effective clearance. This can also be explained by our transplantation experiments with exogenous mucus, where we demonstrated that supplementation of KO cultures with the absent mucin can rescue and improve irregular transport phenotypes. We note supplementation of MUC5B to MUC5B–expressing cultures also improved transport coordination likely by increasing mucin concentration and mucus gel viscoelasticity leading to enhanced hydrodynamic coupling. However overall, these experiments demonstrate the biological importance of both gel-forming mucins, MUC5B and MUC5AC for efficient mucociliary transport in the airways.

Our current understanding of impaired mucus clearance has primarily focused on the imbalance in osmotic pressure between the mucus and periciliary layer, which results from a dehydrated or hyper-concentrated mucus gel that is not easily cleared (*3*, *34*). However, our findings shed light on the mechanisms of defective mucociliary clearance dictated by mucus composition and independent of concentration. While increased MUC5AC may contribute to protection against inhaled pathogens through physical obstruction, disproportionate increases have been associated with pathogenesis of chronic obstructive lung diseases and airway obstruction (*7*, *19*). Mucous plugs from individuals with fatal asthma show MUC5AC enrichment and localization close to the epithelium. In mice, Muc5ac mediates mucous occlusion after allergic stimulation, while Muc5ac-KO resulted in significant protection against mucus plugging. Taken together, elevated levels of MUC5AC on the epithelial surface could be a significant contributor to mucus accumulation and airway obstruction due to its poor transportability as demonstrated in our studies.

In conclusion, our study provides unique insight into the contribution of each gel-forming mucin, MUC5B and MUC5AC on mucociliary clearance mechanisms, demonstrating the significance of each mucin. Our findings suggest secretion of MUC5B and MUC5AC offer a means for the airway epithelium to dynamically control mucus transport velocity and flow alignment. While MUC5AC has been considered to be a potential therapeutic target for allergic asthma (*35*), we predict that preserving baseline levels of MUC5AC could be important in maintaining flow orientation to facilitate effective removal of pathogens. To resolve MUC5AC–induced MCC dysfunction, ongoing work on mucolytic strategies that target the disulfide bonds in mucus gels for normalizing clearance may prove useful as MUC5AC–enriched gels are likely to be more extensively crosslinked (*15*, *36*). This work also establishes the premise for future development of new precision therapies aimed at restoring an optimal balance of MUC5B and MUC5AC secretion in airway diseases.

## Materials and Methods

### CRISPR/Cas9-mediated knockout of MUC5B and MUC5AC

We utilized lentivirus for CRISPR/Cas9-mediated gene targeting of MUC5B and MUC5AC in BCi-NS1.1 cells, an h-TERT immortalized human airway basal cell line, generously provided by Dr. Ronald Crystal, which their group established and characterized in previous work (*25*). Two single guide (sg) RNAs that target different regions of MUC5B and MUC5AC were selected based on favorable targeting using the Doench and Xu scores. To generate negative control, we designed sgRNA sequences with no matching sequence in the genome (non-targeting control). Selected gRNAs were cloned into a GFP-expressing lentiviral vector, pLENTICRISPRv2-GFP, (a gift from Dr. Feng Zhang (Addgene plasmid #52961; http://n2t.net/addgene:52961; RRID:Addgene_52961)), which also encodes Cas9 nuclease. Lentiviral stalks were generated by co-transfecting plentiCRISPRv2, pCMV-VSV-G (a gift from Dr. Bob Weinberg (Addgene plasmid #8454; http://n2t.net/addgene:8454; RRID:Addgene_8454), and psPAX2 (a gift from Dr. Didier Trono (Addgene plasmid #12260; http://n2t.net/addgene:12260; RRID:Addgene_12260)) into HEK293T cells with jetPrime (Polyplus) in cell media, DMEM (Gibco) supplemented with 10% fetal bovine serum using manufacturer’s protocol. BCi-NS1.1 cells were transduced at 40-60% confluence using harvested lentiviruses in media with a final concentration of 20 mM HEPES (Gibco) and 4 μg / mL Polybrene (American Bio). BCi-NS1.1 cells were then centrifuged (1,000 *g* for one hour at 37°C) and incubated at 37°C overnight. At 60-80% confluence cells were passaged and expanded and sorted for eGFP expression.

### T7 endonuclease cleavage assay

DNA from transduced, sorted (eGFP+) cells was extracted using QuickExtract DNA Extraction Solution (Lucigen). EnGen Mutation Detection Kit (New England BioLabs) was used for amplification of target DNA, heteroduplex formation, and heteroduplex digestion, using manufacturer’s protocol. Final DNA products were separated and visualized in a 1% agarose gel through gel electrophoresis.

### Cell Culture

Transduced and sorted (eGFP+) BCi-NS1.1 cells were seeded on plastic at ~3000 cells/cm^2^ in PneumaCult-Ex Plus media (StemCell) and incubated at 37°C, 5% CO_2_. Once cells reached 70-80% confluency, they were dissociated using 0.05% trypsin ethylenediaminetetraacetic acid (EDTA) for 5 minutes at 37°C. To establish well-differentiated human airway epithelial (HAE) tissues cultures grown at an air-liquid interface (ALI), BCi-NS1.1 cells were seeded on 12 mm diameter transwell inserts (Corning Costar) coated with 50 μg/mL type 1-collagen from rat-tail (Corning) at ~10,000 cells/cm^2^. Expansion media, PneumaCult-Ex Plus, was used to feed cells in both the apical and basolateral compartments until 100% confluency. After reaching confluency, the apical media was removed to transition from submerged to ALI culture and the basolateral media was replaced with PneumaCult-ALI (StemCell). All cells were grown for 28 days to reach differentiation with media exchanged every other day. Transepithelial electrical resistance (TEER) was quantified cultures using the Millicell Electrical Resistance System ERS-2 Volt-Ohm Meter (Millipore).

### Mucus collection

Once fully differentiated, mucus was allowed to accumulate prior to collection every 3-4 days. Mucus samples were harvested by washing apical compartments with PBS for 30 minutes at 37°C. After 30 minutes incubation, the solution of mucus and PBS was collected (*11*, *13*). Samples were loaded into Amicon Ultra 100 kDa filters (Millipore-Sigma) and centrifuged at 14,000 *xg* for 20 minutes to remove excess PBS for mucin isolation. Mucus samples were stored at −80 °C for long-term storage (> 2 weeks) or stored at 4 °C (*26*). Pooled mucus samples were used for western blot analysis, solids concentration analysis, biophysical characterization, and transport experiments.

### Western blot analysis

Mucus samples from BCi-NS1.1 cell washings were used for western blot analysis. Protein concentration of samples was quantified using BCA assay (Pierce, Thermo Scientific) and equal amounts (10 μg) were loaded into each lane and run on a 4-20% Tris-Glycine gel (Novex, Invitrogen) under reducing conditions. Proteins were transferred to a polyvinylidene fluoride (PVDF) membrane (GE Healthcare) and blocked with 5% (w/v) fat free milk protein in tris-buffered saline, 0.1% Tween (TBS-T) at room temperature. Primary antibody incubation was over night at 4°C in 5% (w/v) fat free milk protein. The primary antibodies that were used were mouse anti-MU5AC (1:1,000; cat. no. ab3649; Abcam) and rabbit anti-MUC5B UNC414 (1:1,000), generously gifted by Dr. Camille Ehre. After washing in TBS-T, membranes were probed with secondary antibodies for one hour at room temperature in blocking buffer. The secondary antibodies that were used were anti-mouse IgGk-HRP (sc-516102, Santa Cruz, 1:10,000), anti-rabbit-HRP (32460, Invitrogen, 1:10,000). Imaging was performed with chemiluminescent SuperSignal Dura reagent (Thermo Scientific) on an iBright 1500 (Thermo Fisher).

### Mucus solids concentration

We measured solids concentrations of mucus samples by aliquoting 50 μL of mucus on weighing paper of known mass and recording the combined mass of the mucus sample and paper. The samples were then placed on a heat plate at 60°C for 4-5 hours. The final mass of the dried sample and weighing paper was recorded and used to calculate % solids concentration.

### Bulk rheology

Dynamic rheological measurements of mucus gels harvested and pooled from ALI cultures were performed using the ARES G2 rheometer (TA Instruments) with a 40-mm diameter 2° cone and plate geometry at 25°C. To determine the linear viscoelastic region of the fully formed gel, a strain sweep measurement was collected from 0.1-10% strain at a frequency of 1 rad s^-1^. To determine the elastic modulus, *G*′(ω), and viscous modulus, *G*″(ω), a frequency sweep measurement was conducted within the linear viscoelastic region of the gel, at 1% strain amplitude and angular frequencies from 0.1 to 10 rad s^-1^.

### Particle Tracking Microrheology (PTM)

Carboxylate-modified, fluorescent polystyrene nanoparticles (PS-COOH; Life Technologies) with a diameter of 100 nm were coated with a high surface density of polyethylene glycol (PEG) via a carboxyl-amine linkage using 5-kDa methoxy PEG-amine (Creative PEGWorks) as previously reported (*9*, *37*). Particle size and zeta potential was measured in 10 mM NaCl at pH 7 using a NanoBrook Omni (Brookhaven Instruments). We measured diameters of 122 nm and zeta potentials of −0.54 ± 1.16 for 100 nm PEG-coated PS nanoparticles, respectively. The diffusion of the PEG-coated nanoparticles (PEG-NP) in mucus gels was measured using fluorescence video microscopy. Twenty five μL of mucus was added to the chamber along with 1 μL of ~0.002% w/v suspension of PEG-NP 30 minutes prior to particle tracking microrheology (PTM) experiments. Videos were collected using a Zeiss 800 LSM microscope with a 63x water-immersion objective and an Axiocam 702 camera (Zeiss) at a frame rate of 33 Hz for 10 seconds at room temperature. For each sample, at least 3 high-speed videos were recorded. The particle tracking analysis was performed using a previously developed image processing algorithm (*37*, *38*). Mean squared displacement (MSD) as a function of time lag (τ) was calculated as

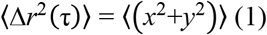

for each particle. Using the generalized Stokes-Einstein relation, measured MSD values were used to compute viscoelastic properties of the hydrogels. The Laplace transform of 〈Δ*r*^2^(*τ*)〉 −6Δ*r*^2^(s)〉, is related to viscoelastic spectrum 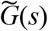 using the equation

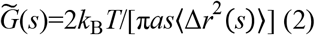

where *k*_B_*T* is the thermal energy, *a* is the particle radius, *s* is the complex Laplace frequency. The complex modulus can be calculated as

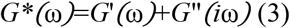

with iω being substituted for *s*, where *i* is a complex number and ω is frequency. Hydrogel network pore size, *ξ*, is estimated based on *G*’ using the equation,

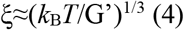

### Mucociliary transport and ciliary beat frequency

For measurement of mucociliary transport rate, a 4 uL suspension of 2 μm red-fluorescent polystyrene microspheres (Sigma-Aldrich; 1:1000 dilution in PBS) was apically applied to native mucus, which was allowed to accumulate for 1-2 days prior to analysis. After equilibration overnight, videos of three regions were recorded at 10x magnification using a Zeiss 800 LSM microscope. Images were collected at a frame rate of 0.5 Hz for 20 seconds on the plane of the mucus gel. The microsphere tracking data analysis is based on an image processing algorithm that was custom written in MATLAB (*26*, *39*). Briefly, the analysis software computes the xy-plane trajectories of each fluorescent microsphere in each frame. Using the trajectory data, displacement of microspheres was computed, and transport rate was calculated by dividing the distance traveled by total elapsed time. In order to measure ciliary beat frequency (CBF), 10 second videos at a frame rate of 20 Hz were recorded at 20x magnification in ≥3 randomly selected regions of each BCi-NS1.1 culture using brightfield. Using a custom written algorithm in MATLAB, the number of local pixel intensity maxima were counted, which indicates beating of cilia. Beat frequency was determined by dividing the number of beats over the total elapsed time.

### Immunostaining

BCi-NS1.1 cultures were fixed using 4% paraformaldehyde for 30 minutes at room temperature. Inserts were washed with PBS and blocked with 5% bovine serum albumin (BSA) in PBS and 0.01% triton X-100 for 1 hour at room temperature. After blocking, fixed cultures were incubated overnight at 4°C with AlexaFluor conjugated antibodies in 5% BSA in PBS. The antibodies that were used were anti-acetylated alpha tubulin AlexaFluor 647 (cilia marker, clone 6-11B-1, sc23950, Santa Cruz, 1:2000) and ZO-1 monocolonal antibody AlexaFluor 488 (tight junction marker, ZO1-1A12, Invitrogen, 1:2000). Inserts were washed with PBS three times and imaged at 63x magnification using a Zeiss 800 LSM microscope (Zeiss). Quantification of % tubulin positive area was performed in FIJI using the automated thresholding function.

### Mucus Transplantation

Reconstituted mucus samples from KO culture washings were used for transplantation studies. 20 μL of mucus was applied to the apical surface of ALI cultures that had been washed with PBS for 30 minutes (*26*, *40*). 2 μm red-fluorescent polystyrene microspheres were applied immediately after transplantation of exogenous mucus. Mucociliary transport rates were measured after equilibration overnight as described in the methods above.

### Statistical Analysis

All graphing and statistical analyses were performed using GraphPad Prism 8 (GraphPad Software). Two-group comparisons were performed using 2-tailed Student’s *t*-test (normally distributed data) or Mann-Whitney *U* test. For comparisons between groups, one-way analysis of variance (ANOVA) followed by a Tukey *post hoc* correction was performed. Kruskal-Wallis with Dunn’s correction was used for comparison of multiple groups with non-Gaussian distributions. Bar graphs show mean and standard deviation and box and whiskers plots show median. Differences were considered statistically different at the level of *p*< 0.05.

## Acknowledgments

We acknowledge the BioWorkshop core facility in the Fischell Department of Bioengineering at the University of Maryland – College Park for use of their dynamic light scattering instrument, microplate reader, and rheometer.

## Author contributions

Conceptualization: DS, MAS, GAD
Methodology: DS, EI, MAS, GAD
Investigation: DS, EI
Visualization: DS, LK
Supervision: MAS, GAD
Writing—original draft: DS
Writing—review & editing: DS, EI, MAS, GAD

## Data and materials availability

All data are available in the main text or the supplementary materials.

