## Supplementary Materials for "MUC5B mobilizes and MUC5AC spatially aligns mucociliary transport on human airway epithelium"

**Table.S1. sgRNA sequence and target**

| Gene | gRNA # | Designation | Target | Sequence |
|---|---|---|---|---|
| MUC5B | gRNA 1 | B KO1 | Exon 3 | GTGGAACAAAGCTCACGCGC |
| | gRNA 2 | B KO2 | Exon 4 | TTCAACGTCCAGCTACGCCG |
| MUC5AC | gRNA 1 | AC KO1 | Exon 4 | GATGTTAAAATCCTCGTAGG |
| | gRNA 2 | AC KO2 | Exon 6 | GAGAGGAGCTCGCTGACCAC |

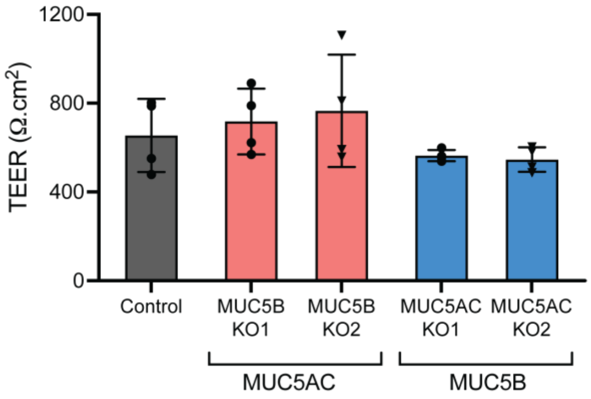


**Fig. S1. Transepithelial electrical resistance in differentiated control and MUC5B/MUC5AC KO BCi-NS1.1 HAE cultures.**

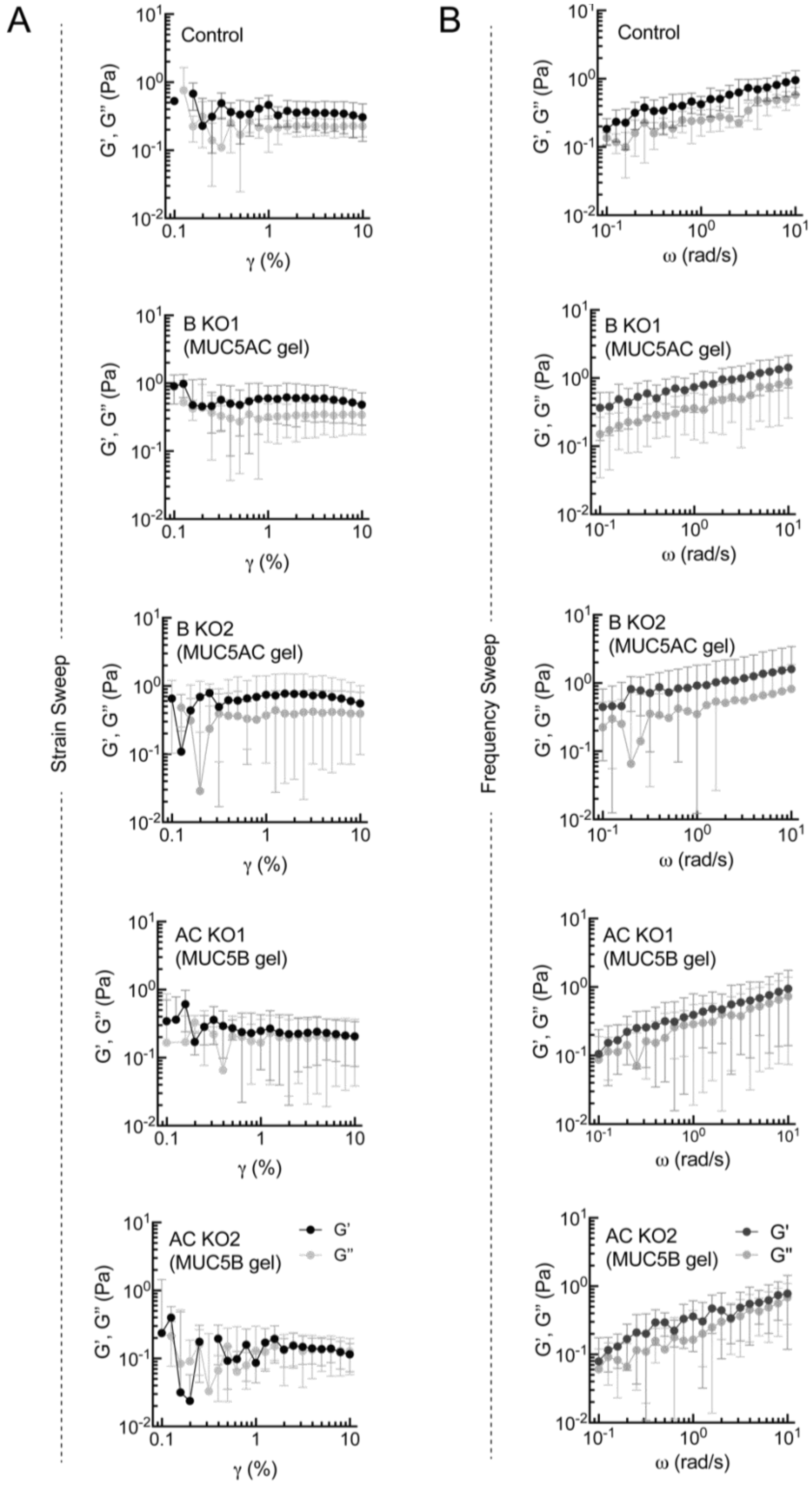


**Fig. S2. Viscoelastic properties of mucus produced from MUC5B/AC-KO cultures. (A)** strain sweep at ω = 1 rad/s and **(B)** frequency sweep at 1% strain in mucus produced from MUC5B/AC-KO cultures.

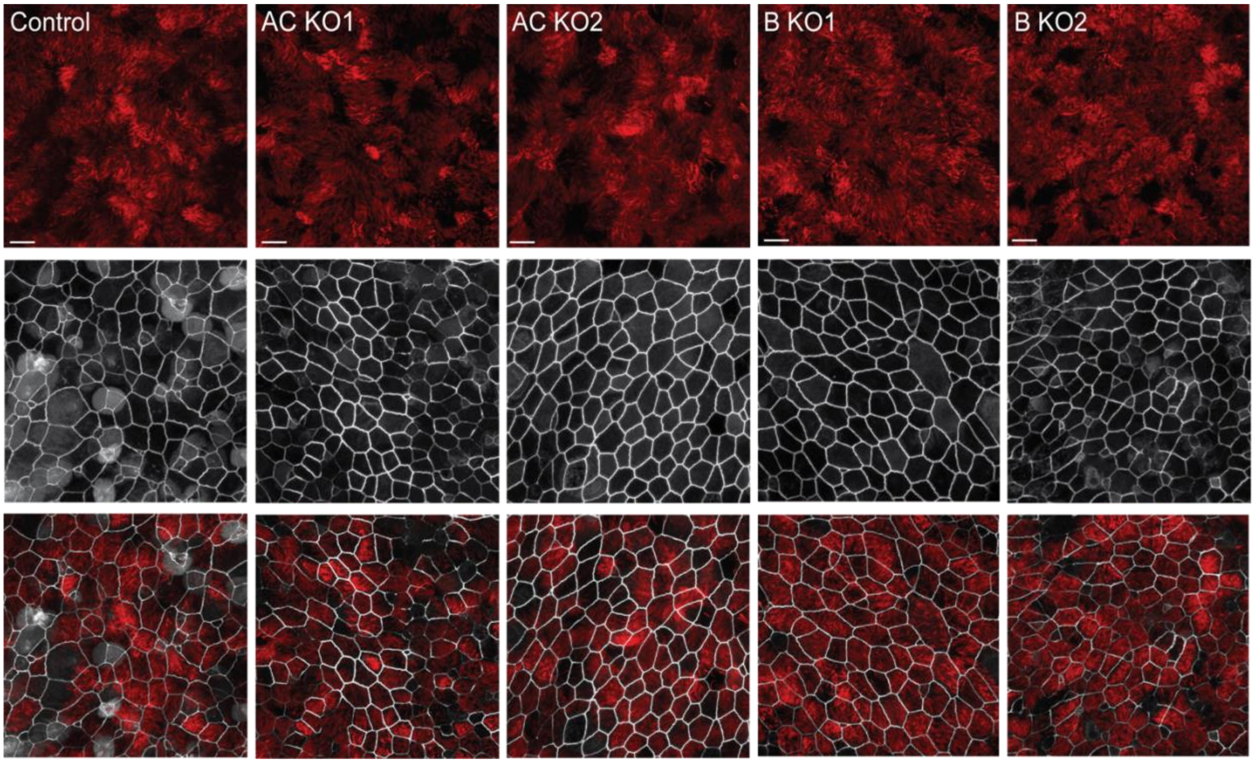


**Fig. S3. Representative images of differentiated MUC5B/AC-KO HAE cultures with immunostaining for acetylated alpha tubulin (red) and ZO-1 tight junction associated protein (white).**
